# KIAA0319 influences cilia length, cell migration and mechanical cell-substrate interaction

**DOI:** 10.1101/780874

**Authors:** Rebeca Diaz, Nils M. Kronenberg, Angela Martinelli, Philipp Liehm, Andrew C. Riches, Malte C. Gather, Silvia Paracchini

## Abstract

Following its association with dyslexia in multiple genetic studies, the KIAA0319 gene has been extensively investigated in different animal models but its function in neurodevelopment remains poorly understood.

We developed the first cellular knockout model for KIAA0319 via CRISPR-Cas9n to investigate its role in processes suggested but not confirmed in previous studies, including cilia formation and cell migration. We found that KIAA0319 knockout increased cilia length and accelerated cell migration. Using Elastic Resonator Interference Stress Microscopy (ERISM), we detected an increase in cellular force for the knockout cells that was restored by a rescue experiment. Combining ERISM and immunostaining we show that KIAA0319 depletion reduces the number of podosomes formed by the cells.

Our results suggest an involvement of KIAA0319 in cilia biology and force regulation and show for the first time that podosomes exert highly dynamic, piconewton vertical forces in epithelial cells.

## Introduction

Dyslexia is a neurodevelopmental disorder that affects around 5% of school-aged children and refers to unexpected difficulties in learning to read [1]. Dyslexia is high heritable (up to 70%). Genetic studies, mainly in family-based samples, have focused their attention on genes that play a role in neurodevelopment, including *DYX1C1, DCDC2, ROBO1* and *KIAA0319* [2]. Functional analysis of these genes have largely contributed to shape hypotheses aimed at explaining the neurobiology of dyslexia. Initial *in utero* gene silencing experiments in rats for these genes provided strong support for the neuronal migration hypothesis [3] first proposed in the eighties [4]. This hypothesis is based on the observation of subtle cortical anomalies, i.e. heterotopias and microgyrias, in post-mortem brains from individuals with dyslexia (*n* = 8). Such anomalies are likely to be the result of neuronal migration defects. However, knockout mouse models for DYX1C1, DCDC2 and KIAA0319 did not exhibit cortical alterations [5], although heterotopias were observed in rats subjected to *in utero* knockdown of Dyx1c1 [6]. The discordance between knockdown experiments in rat and knockout mouse models has been explained by species-specific effects, compensatory mechanisms in mice, or artefacts in shRNA experiments, and has been highlighted by recent reviews of the literature, providing interpretations either in support or raising doubts about the neuronal migration hypothesis [5,7,8].

In parallel, emerging evidence supports roles of *DCDC2, DYX1C1, ROBO1* and *KIAA0319* in cilia biology [2]. Transcriptomic studies showed differential expression for these genes in ciliated tissue [9–11]. Beyond these studies, a role of KIAA0319 in cilia biology has not been described yet, but cellular and animal knockouts for DCDC2 and DYX1C1 presented cilia defects. Mutations in DYX1C1 and DCDC2 have been identified in patients with ciliopathies, a group of disorders caused by defective cilia and often characterised by alterations in body asymmetry. ROBO1 has been shown to localize to the cilium of mouse embryonic interneurons. Cilia biology has been proposed as a molecular link to explain the atypical brain asymmetries which are consistently reported for neurodevelopmental disorders, such as dyslexia [12,13].

*KIAA0319* encodes a transmembrane protein with five PKD domains [14,15] (Figure 1A). Such structures have been previously found in cell surface proteins and are known to be involved in cell-cell and cell-matrix interactions [16] but the cellular function of KIAA0319 remains largely uncharacterised [5]. KIAA0319 has also been suggested to play a role in signalling pathways [17] and in axon growth inhibition [18]. A gene expression analysis in zebrafish showed very high expression in the first hours of development and specific signal in defined embryonic structures, including the notochord and the developing eye and otic vesicles [19]. Moreover, in support of the importance of KIAA0319 in brain function, a recent study suggests a possible role of KIAA0319 in Alzheimer’s disease [20].

**Figure 1.**
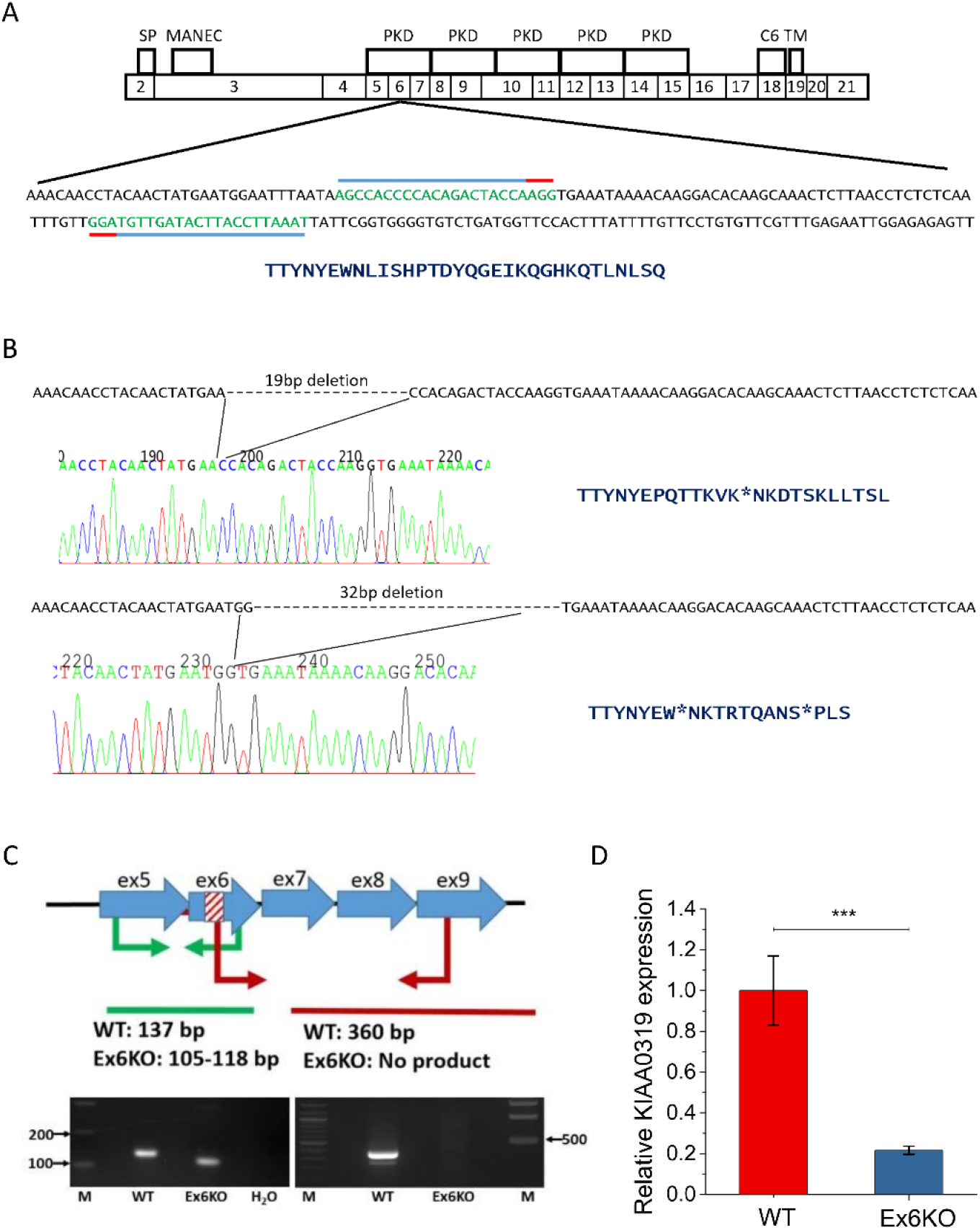
Generation of a cellular KIAA0319 knockout. **(A)** Top: Structure of Human KIAA0319 (based on [15] and Ensembl release 94 [51]). The diagram shows the correspondence between protein domains and coding exons in KIAA0319. Signal peptide (SP), MANEC domain (MANEC), PKD domains (PKD), cysteine residues (C6) and transmembrane domain (TM) are indicated. Bottom: full DNA sequence of KIAA0319 exon 6 with target sequences for the gRNAs indicated with blue lines. Red lines show the position of the PAM sequences. Translated sequence of amino acids for the targeted exon is shown below the DNA sequence. **(B)** Chromatograms of the deletions found in Ex6KO and translated corresponding amino acids for wild type and knockout cell line. Asterisks indicate premature stop codons. **(C)** Results of the PCR screening to confirm the deletions in Ex6KO. The cartoon on the left represents the screening strategy. Two sets of primers were designed to give different bands in the WT and KO. The stripped area indicates the 19 and 32 base pair (bp) deletions in the exon 6 of KIAA0319. The first set of primers (Ex_6R and Ex_5F) amplifies the region around the deletion and therefore a smaller band is expected for the KO (105 – 118 bp) compared to the WT (137 bp). The second pair (Ex9_R/Ex6delF) has one primer mapping within the deletion. PCR is expected to give a band of 360 bp in the WT and no product in the KO. Images below confirm the expected results for both pairs. **(D)** Quantification of KIAA0319 mRNA in WT and Ex6KO by qPCR. KIAA0319 expression is significantly lower in Ex6KO (Student’s *t*-test: *p* ≤ 0.0001), consistent with nonsense mediated decay of the mRNA caused by the deletion.

In spite of these intensive efforts, the precise role of KIAA0319 remains unexplained. The initial neuronal migration hypothesis is not consistently supported, and direct evidence for a role in cilia biology, similarly to other genes implicated in dyslexia, has not been described yet. To address this issue, we generated the first cellular knockout model of KIAA0319 in human cells to specifically investigate its role in cilia biology and neuronal migration, addressing the two main hypotheses currently proposed. We used retina pigmented epithelium cells (RPE1), which are particularly suitable to study cilia, and studied their mechanobiology using a range of assays including the recently introduced Elastic Resonator Interference Stress Microscopy (ERISM) [21,22] that allows for continuous imaging of cell forces with high spatial resolution and over extended periods of time.

We show that loss of KIAA0319 leads to longer cilia, changed migratory behaviour and increased cellular force exertion. In addition, our data indicate that KIAA0319 knockout cells form fewer podosomes, structures that have been shown to have mechanosensitive function via the exertion of oscillating, vertical forces [23]. However, mechanical activity of podosomes had not been observed and measured in epithelial cells before.

## Results

### Generation of KIAA0319 KO in RPE1 cell lines

We generated KIAA0319 knockout RPE1 cells with CRISPR-Cas9n based genome editing. The *KIAA0319* main isoform (NM_014809) consists of 21 exons and spans 102 kb of human chromosome 6 (Figure 1A). We generated a biallelic knockout (Ex6KO) by causing deletions that introduce premature stop codons at exon 6 of *KIAA0319* using paired gRNAs (Figure 1B). The deletion was confirmed by RT-PCR (Figure 1C). Transcript quantification by qRT-PCR shows that KIAA0319 expression in Ex6KO is five-times lower than the wild-type (*t*-test, *p* ≤ 0.001) which is consistent with degradation of the transcript by nonsense-mediated decay[24] (Figure 1D). We characterised the KIAA0319 knockout to address two specific hypotheses emerging from the literature. Specifically we tested whether KIAA0319 might play a role in i) cilia and ii) neuronal migration.

### KIAA0319 knockout cells form longer primary cilia

We measured cilia length in RPE1 wild type (WT) and Ex6KO cells by staining of the cilium-specific protein ARL13B and analysis of epi-fluorescence images (Figure 2A & B). While a similar fraction of WT and Ex6KO cells formed cilia (WT: 379/571, 68%; Ex6KO: 271/383, 70%), the cilia in Ex6KO were significantly longer than in the wild type (mean ± SEM: WT: 4.5 µm ± 0.1 µm, *n* = 129; Ex6KO: 6.1 µm ± 0.2 µm, *n* = 104; *t*-test: *p* ≤ 0.001; Figure 2C).

**Figure 2.**
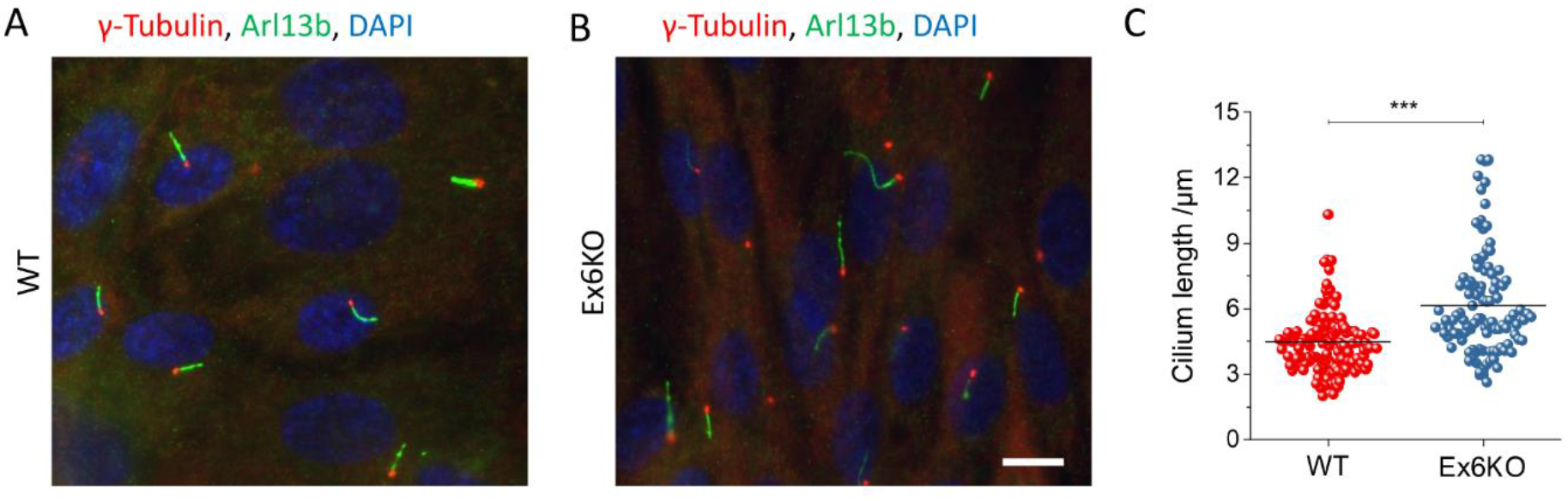
Analysis of the cilia length. Representative immunofluorescence images of RPE1 wild type (**A**) and Ex6KO (**B**), stained for cilia marker Arl13b (green), centrosomal marker γ-tubulin (red), and DAPI (blue). **(C)** Plot of the cilia length for wild type (*n* = 129) and Ex6KO cells (*n* = 104). Groups were compared using the Student’s *t*-test (‪‪‪: *p* ≤ 0.001). Scale bar, 10 µm.

### Migration speed, cell morphology, and force exertion are altered in KIAA0319 knockout cells

The second hypothesis we investigated was the role of KIAA0319 in cell migration. We started by comparing WT and KO with a scratch assay on confluent layers of cells to test collective cell migration. The assays did not reveal a significant difference in the capacity to cover the empty space between WT and Ex6KO cells after 24 h (mean cell coverage ± SEM: WT: 27.4% ± 4.2%, *n* = 3; Ex6KO: 30.2% ± 3.5%, *n* = 3; *t*-test: *p* = 0.63; Figure S1).

Next, we turned to investigations on the single cell level, characterizing migration speed and cell morphology through detailed analysis of phase contrast microscopy and, in parallel, mapping the mechanical forces exerted by the cells using ERISM. For ERISM, cells are cultured on substrates that consist of a layer of an ultra-soft elastomer situated between two semi-transparent, mechanically flexible gold mirrors, which form an optical micro-cavity. Mechanical force and stress exerted by cells cause local deformations of the soft micro-cavity (the effective stiffness of cavities used in this study is 6 kPa). This in turn leads to local shifts in cavity resonance that are analysed by optical modelling to compute a high-resolution displacement map with µm lateral resolution and nm vertical resolution [22], which allows for the detection of forces in the piconewton range. In our earlier work [21], ERISM has been extensively calibrated (through application of well-defined forces by an atomic force microscope) and validated (through measurements on widely studied cell lines). The substrates used for ERISM are semi-transparent, thus allowing to combine force mapping with high-resolution imaging.

WT and Ex6KO cells were plated on ERISM substrates at a density low enough to ensure non-confluency and thus allow the behaviour of individual cells to be analysed (Figure 3A).Quantitative image analysis showed that Ex6KO cells covered a smaller surface area than WT cells (mean cell area ± SEM: WT: 2052 µm^2^ ± 91 µm^2^, *n* = 36; Ex6KO: 1295 µm^2^ ± 65 µm^2^, *n* = 36; *t*-test: *p* ≤ 0.001; Figure 3B), even though the shape and morphology of the cells did not differ. The displacement maps recorded with ERISM (Figure 3A) revealed that cells from both lines generated similar spatial patterns of force on their substrate (even though absolute forces were markedly different, see next section): pulling was focused around the two long ends of the cells, perpendicular to the direction of migration (cells were polarised in a way that the nucleus was positioned posterior to the direction of migration). Downward compression was observed underneath the centre of the cells. This displacement pattern is a fingerprint for the exertion of contractile forces by adherent cells [21].

**Figure 3.**
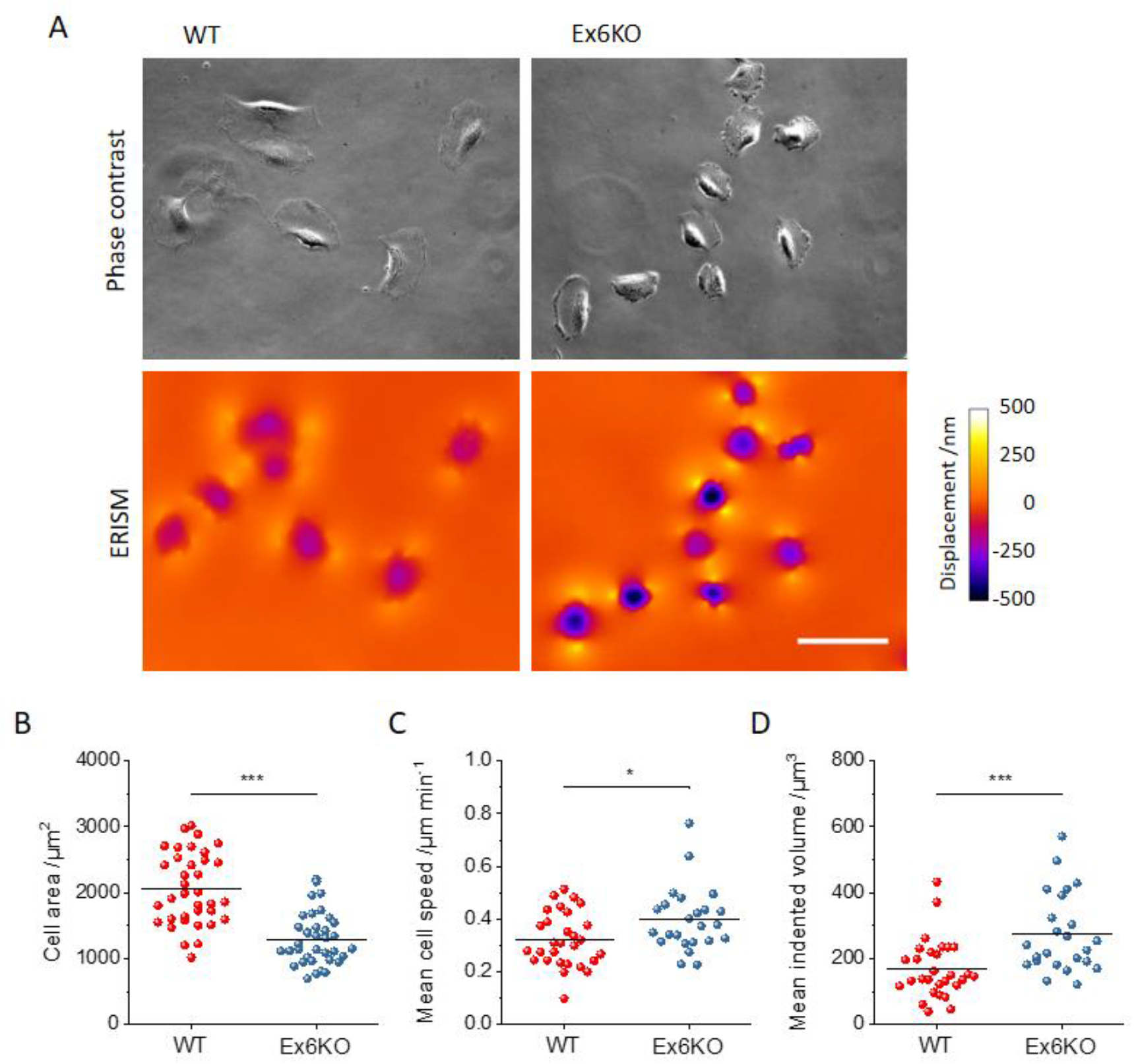
Analysis of mechanical activity of RPE1 WT and Ex6KO cells during migration on an ERISM micro-cavity. **(A)** Phase contrast (upper row) and ERISM micro-cavity displacement maps (lower row) of WT (left) and Ex6KO (right) cells. **(B)** Comparison of the surface area covered by WT (*n* = 36) and Ex6KO (*n* = 36) cells types. **(C)** Comparison of mean speed of WT (*n* = 29) and Ex6KO (*n* = 24) cells. **(D)** Comparison of mean indented volume of WT (*n* = 29) and Ex6KO (*n* = 24) cells. Only cells with free movement for ≥2 h were included in analysis of speed and indented volume. Plots in (B), (D) and (E) show all measured data points and the mean (line). Groups were compared using the Student’s *t*-test (‪: *p* ≤ 0.05, ‪‪: *p* ≤ 0.01, ‪‪‪: *p* ≤ 0.001). Scale bar, 50 µm.

The migratory behaviour and the associated dynamics of force exertion of WT and Ex6KO cells were then investigated by taking time-lapse measurements of phase contrast and ERISM displacement maps in five-minute intervals over a time span of 17 hours (Movie S1 & S2). The average speed of single cell migration was significantly higher for Ex6KO than for WT cells (mean speed ± SEM: WT: 0.32 µm min^-1^ ± 0.02 µm min^-1^, *n* = 29; Ex6KO: 0.40 µm min^- 1^ ± 0.02 µm min^-1^, *n* = 24, *t*-test: *p* = 0.02; only cells with free movement for ≥2 h were included in analysis; Figure 3C). To assess the force exerted by cells, we computed the total volume by which each cell indents into the substrate and used this as a proxy for the applied force [21]. Comparing the temporal averages of applied force during migration showed that Ex6KO cells exerted significantly stronger contractile forces on the substrate than WT cells (mean indented volume ± SEM: WT: 168 µm^3^ ± 16 µm^3^, *n* = 29; Ex6KO: 273 µm^3^ ± 24 µm^3^, *n* = 24; *t*-test: *p* = 0.0006; only cells with free movement for ≥2 h were included in analysis; Figure 3D).

Next, we investigated changes in migration speed and exerted force over time (Figure S2A). WT and Ex6KO cells showed no differences in how migration speed and applied force fluctuated when normalising data to the respective means (Figure S2B & C). For both WT and Ex6KO cells, intervals of increased migration speed correlated with drops in cell forces (anti-correlation between the first time derivative of speed and the first time derivative of mechanical activity). Again, there was no significant difference in this correlation between the two groups (Figure S2D, F & G). Furthermore, the straightness of the migration was not affected by the KIAA0319 knockout (Figure S2H).

To validate our findings of the impact of KIAA0319 on cell area and cell force, we conducted a rescue experiment by generating an Ex6KO cell line with stable expression of KIAA0319-GFP fusion protein (Ex6KO K-GFP; Figure 4A). We also generated a control line of RPE1 WT cells with the same construct (WT K-GFP). Even though the KIAA0319 rescue did not recover the reduction in cell area seen for Ex6KO cells [mean cell area ± SEM: WT: 2315 µm^2^ ± 200 µm^2^, *n* = 16; WT K-GFP: 2299 µm^2^ ± 107 µm^2^, *n* = 20; Ex6KO: 1565 µm^2^ ± 123 µm^2^, *n* = 23; Ex6KO K-GFP: 1297 µm^2^ ± 131 µm^2^, *n* = 17; Figure 4B], the level of cell force was restored in Ex6KO K-GFP cells [mean indented volume ± SEM: WT: 115 µm^3^ ± 14 µm^3^, *n* = 16; WT K-GFP: 96 ± 9 µm^3^, *n* = 19; Ex6KO: 186 ± 20 µm^3^, *n* = 24; Ex6KO K-GFP: 125 ± 16 µm^3^, *n* = 16; *t*-test(WT vs. Ex6KO): *p* = 0.01, *t*-test(WT vs. Ex6KO K-GFP): *p* = 0.67; Figure 4C)].

**Figure 4.**
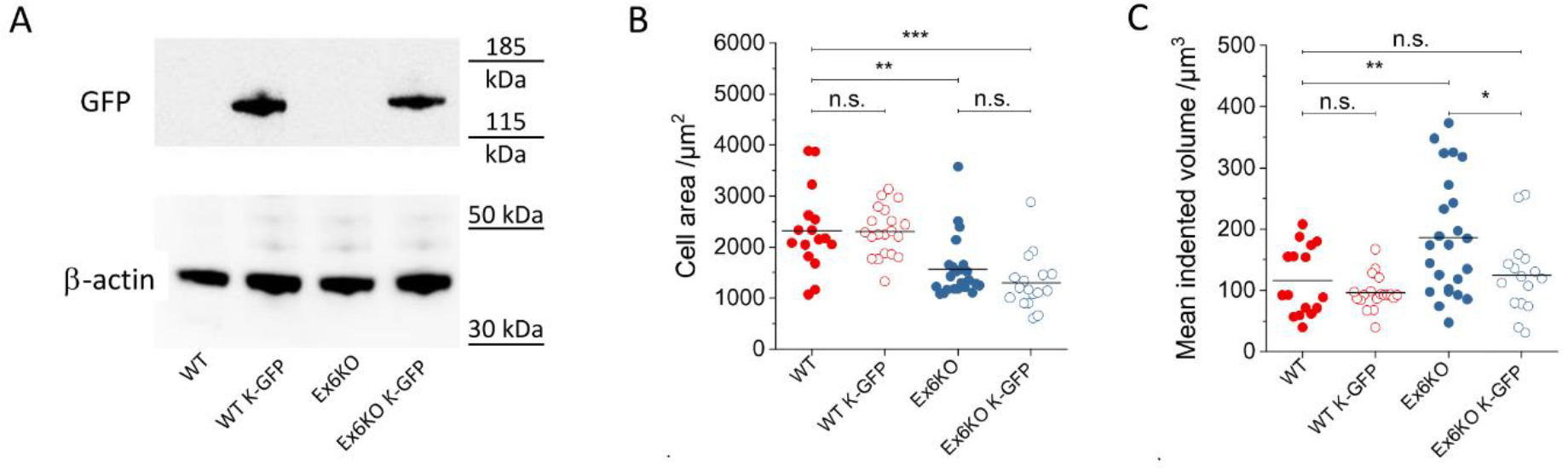
Phenotype recovery through KIAA0319 rescue. **(A)** Western blot confirming the presence of a fusion protein (140 KDa) following transfection with a full length KIAA0319 construct fused to a GFP tag. **(B)** Comparison of area covered by RPE1 WT, WT K-GFP, Ex6KO and Ex6KO K-GFP cells attached to ERISM micro-cavity. (WT: *n* = 16, WT K-GFP: *n* = 20, Ex6KO: *n* = 23, Ex6KO K-GFP: *n* = 17) **(C)** Comparison of mean mechanical activity of RPE1 WT, WT K-GFP, Ex6KO and Ex6KO K-GFP cells during migration on ERISM micro-cavity. Only cells with free movement for >4 h were included in the analysis. Plots in B and C show measured data points and the mean (line). (WT: *n* = 16, WT K-GFP: *n* = 19, Ex6KO: *n* = 24, Ex6KO K-GFP: *n* = 16) Groups were compared using the Student’s *t*-test (‪: *p* ≤ 0.05, ‪‪: *p* ≤ 0.01, ‪‪‪ *p* ≤ 0.001).

### RPE1 KIAA0319 WT and Ex6KO show different fine patterns of force exertion

Given the differences in cilia length, cell area, migration speed and exerted force, we reasoned that KIAA0319 knockout might affect cytoskeleton dynamics. To test this hypothesis, we took phase contrast and ERISM time-lapse measurements of migrating WT and Ex6KO cells at 5 seconds intervals (Movie S3 & S4), and fixed and immunostained the cells for actin and vinculin immediately after the time-lapse.

For further analysis, spatial Fourier-filtering of ERISM maps was used to filter out broad deformation features associated with the overall contractility of cells and thus resolve finer details linked to interaction of sub-cellular components, e.g. focal adhesions or podosomes, with the substrate [21]. (For further discussion on the displacement fine-structure in Fourier-filtered displacement maps see Figure S3.) Figure 5A shows phase contrast images, Fourier-filtered ERISM maps and immunofluorescence microscopy images for a WT and Ex6KO cell. The Fourier-filtered displacement maps of both cells showed numerous small push-pull features that co-localised with vinculin-rich areas in the immunofluorescence microscopy images (see insets ii and iii to Figure 5A for examples of this feature). Vinculin is enriched in the centre between pulling (red areas in Fourier-filtered ERISM maps) and pushing (green areas). The actin fibres are connected to vinculin. Push-pull features in Fourier-filtered ERISM maps were previously attributed to focal adhesions transmitting horizontal forces that are generated by the actin cytoskeleton to the substrate [21]. In agreement with these earlier observations, the axes defined by the push-pull features co-aligned with the actin fibres that connect different vinculin-rich sites (best visible in the Ex6KO cell in Figure 5A and Figure S3). This push-pull behaviour is also consistent with earlier observations of torque being applied by focal adhesions [25].

**Figure 5.**
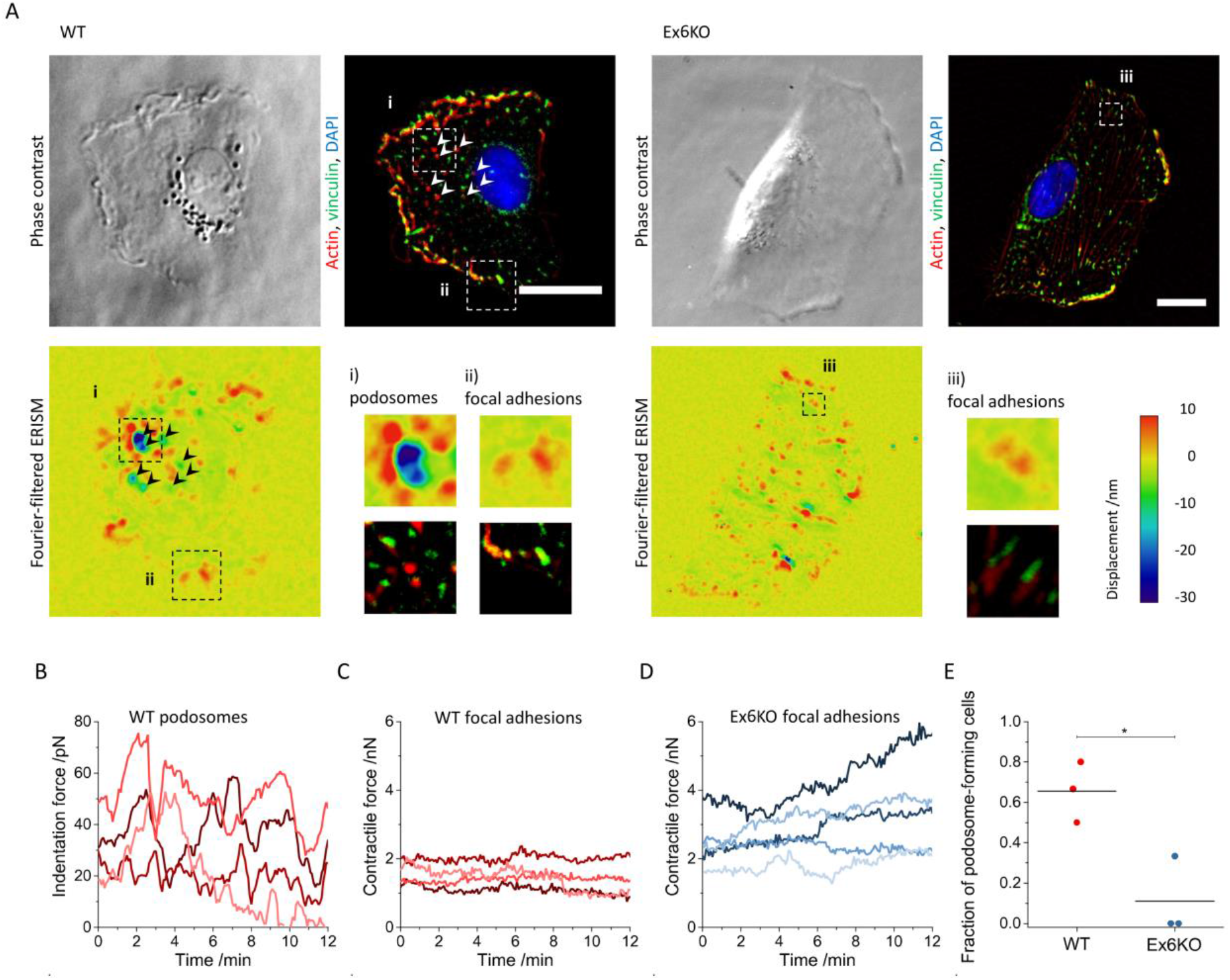
RPE1 KIAA0319 WT and Ex6KO cells use different modes of force exertion. **(A)** Phase contrast images (upper row), Fourier-filtered ERISM displacement maps (middle row) and epi-fluorescence images (lower row, red: actin, green: vinculin, blue: nuclear DNA) of a RPE1 WT cell (left column) and an Ex6KO cell (right column). Arrow heads indicate positions of actin-rich cell protrusions (podosomes). The insets i) in the Fourier-filtered ERISM map and the epi-fluorescence image of the WT cell show magnifications of podosome protrusions. The insets ii) and iii) show magnifications of vinculin-rich cell-substrate contacts (focal adhesions) for the WT and Ex6KO cell, respectively **(B)** Temporal evolution of the indentation force applied by different podosomes of the WT cell shown in A. **(C)** and **(D)** Temporal evolution of the contraction force applied by different focal adhesions of the WT and Ex6KO cell shown in A, respectively. **(E)** Comparison of the fraction of podosome-forming WT and Ex6KO cells. Each data point represents the mean value of an independent experiment investigating *n* = 5 (80%), 3 (67%) and 4 (50%) WT and *n* = 5 (0%), 6 (0%) and 9 (33%) Ex6KO cells, respectively. The lines depict the means. Groups were compared using the Student’s t-test (‪: *p* ≤ 0.05). All scale bars: 20 µm.

The formation and alignment of stress fibres was less distinct in the WT cell than the Ex6KO cell. As a result, the above-mentioned co-alignment of actin, vinculin and ERISM push-pull features was also less pronounced for the WT cell. In agreement with this, the forces exerted by single focal adhesions were smaller in the WT cell than in the Ex6KO cell (Figure 5C & D).

Besides the push-pull features associated with focal adhesions, the Fourier-filtered ERISM displacement maps also showed tightly confined pushing sites with a diameter of about 2 µm (best visible as green-blue areas highlighted with black arrow heads in Fourier-filtered ERISM map of Figure 5A; see inset i to Figure 5A for an example of this feature). These pushing sites were surrounded by circularly arranged dots of upward pulling (red areas). Immunocytochemistry analysis showed that the pushing sites corresponded to actin-rich locations (white arrow heads in epi-fluorescence image of Figure 5A), whereas pulling around the pushing sites colocalised with vinculin-rich positions (inset i to Figure 5A). This protein arrangement is a hallmark of podosomes, a cellular adhesion structure that is chiefly formed in monocyte-derived cells [26] but that has also been reported in spreading and migrating epithelia cells [27].

The time-lapse measurement revealed that the podosomes exerted an oscillating vertical force that reached maximum values of up to 80 pN (Figure 5B). The horizontal contractile forces transmitted by focal adhesions were roughly 100-times larger than the vertical indentation forces exerted by podosomes (Figure 5C & D). However, while podosomal pushing was highly dynamic, the horizontal forces originating from focal adhesions were relatively static and showed little oscillation in force. Focal adhesions at the leading edge of the cell were chiefly stationary once assembled (top right in Movie S4) and any lateral movement of focal adhesions was confined to the trailing edge of the cell (bottom left in Movie S4).

The WT and Ex6KO cell shown in Figure 5 and Movie S3 & S4 are examples illustrating the general differences between the two force transmission patterns (podosomes and focal adhesion). In total, combined ERISM and immunochemistry measurements were carried out for 32 cells in three independent experiments (see Figure S4 and Movie S5 & S6 for further examples). While both WT and Ex6KO cells formed podosomes, we found that the number of cells with podosomes were significantly lower for Ex6KO than for WT cells (fraction of podosome-forming cells ± SEM: WT: 0.66 ± 0.09, *n* = 5, 3 and 4; Ex6KO: 0.11 ± 0.11, *n* = 5,6 and 9, *t*-test: *p* = 0.02).

## Discussion

We successfully developed a cellular knockout model via CRISPR-Cas9n to study the potential role of the *KIAA0319* gene in cilia and cell migration on the basis of the proposed roles of this gene in the literature [2]. Overall, we found that loss of KIAA0319 in RPE1 cells results in elongation of the cilia and an increase of the force the cells apply on the substrate.

Our results suggest a role of KIAA0319 as negative regulator of cilia length. Although the same fraction of KIAA0319 knockout and WT cells developed cilia, these were significantly longer in the knockout (Figure 2C). Atypical cilia length has also been described for the knockdown of Nuclear Distribution Element 1 (*NDE1*), another gene associated with brain cortex development [28]. Cilia formation is a tightly regulated process, and cilia length has functional consequences on processes that include cell cycle re-entry and left-right patterning. Cilia biology is emerging as a contributing factor to a range of diseases, including neurodevelopmental disorders [2]. Other genes implicated in dyslexia have been reported to affect cilia length as well. Knockouts for dyx1c1 present shorter cilia than the wild type in zebrafish [29], and overexpression of Dcdc2 increases cilia length in rat neurons [30]. The only previous evidence in support of a role of KIAA0319 in cilia comes from transcriptomic studies [9–11]. Our work is therefore the first study to support a role for KIAA0319 in cilia biology in a biological model and paves the way for future studies aimed at dissecting the cellular function of this protein.

Investigations on soft ERISM substrates showed that KIAA0319 knockout cells move significantly faster than wild type cells (*p* = 0.02; Figure 3C) and suggest, that KIAA0319 plays a role in regulating single cell migration. However, when assessing collective cell migration with the commonly used scratch assay, we did not observe a significant effect of the KIAA0319 knockout (Figure S1). Several factors might contribute to the different results obtained in these two experiments. First, individual cells might have different migration properties than a layer of collectively migrating cells. Second, the apparent stiffness of the ERISM sensor is in the range of soft tissue (1 to 20 kPa) and significantly different from the stiffness of the cell culture plastic plate in which the scratch assay was performed (∼100,000 kPa) [31]. Substrate stiffness has a strong influence on cell migration in vitro [32]. Furthermore, while cells respond to an acute event, namely local damage, in the scratch assay, the ERISM assay observes the migration of undisturbed cells. Cell attachment proteins are another important factor in cell migration *in vitro*. Both the scratch assay and the ERISM measurements initially used serum containing media and we thus expect that proteins in the serum adhere to the substrate in both cases. However, in the case of the scratch assay, the medium was changed to serum free media after performing the scratch.

The ERISM analysis further revealed that the knockout cells exert significantly stronger forces on their substrate compared to the wild type (Figure 3D). A rescue experiment recovered the mechanical activity of the wild type phenotype (Figure 4C), supporting an involvement of KIAA0319 in cellular forces.

Fluorescent staining strongly indicated the presence of podosomes in both WT and KIAA0319 knockout cells (Figure 5). By combining fluorescence staining with Fourier-filtered ERISM measurements, we found that the actin cores of podosomes protruded vertically into the substrate, exerting oscillating forces of up to 80 pN, while surrounding rings of pulling sites were tightly colocalised with vinculin. Our work shows for the first time that epithelial podosomes mechanically probe the environment by exerting oscillating forces in the pN-range, similarly to what has been previously described for podosomes formed by macrophages [21,23]. While both WT and Ex6KO cells formed podosomes, we found that the fraction of cells with podosomes were significantly lower for KIAA0319 knockout than for WT cells (Fig 5E).

KIAA0319 is a transmembrane protein that contains five PKD domains. These domains have been described in very few human proteins, the best characterised of which is Polycystin-1 (PC1). PC1 acts as a mechanosensor in the membrane of cilia [33], most probably by unfolding of the highly extensible PKD domains in response to stretching forces. It has been proposed that this unfolding maintains contact between neighbouring cells during cell movement [34]. PC1 interacts with the cytoskeleton [35] and plays an important role in adaptative cilia shortening (for example under strong flow) [36]. Therefore, our results suggest that KIAA0319 has a similar function to PC1, affecting both cilia formation and mechanosensing. The knockout of KIAA0319 not only results in formation of longer cilia, but also in an upregulation of mechanical forces. The reduction of podosome formation in KIAA0319 knockout cells compared to the WT further supports the involvement of KIAA0319 in cellular mechanosensing, as the function of podosomes ranges from cell-matrix adhesion and matrix degradation to mechanosensing [26]. In epithelial cells, podosomes were found to associate with hemidesmosomes [27], adhesive structures specific to epithelial cells that regulate a wide range of biological processes including, among others, cell migration, exertion of traction force and mechanosensing [27,37–40].

A difficulty in studying KIAA0319 is the lack of a specific antibody able to detect endogenous levels of this protein. We overcame this by validating the knockout with a combination of approaches: we confirmed loss-of-function deletions in the sixth exon of KIAA0319 that cause stop codons early in the transcript (exon 6 out of 21), so that the knockout cells cannot produce a functional protein. We detected a strong decrease in the expression of KIAA0319, consistent with nonsense mediated decay of the transcript (Figure 1D) due to the loss-of-function deletions.

In summary, our work shows that knockout of KIAA0319 affects processes controlled by the cytoskeleton including cell migration, cilia length, podosome formation and cellular forces. Earlier studies showed that KIAA0319 overexpression inhibits axon growth and KIAA0319 knockout results in neurite outgrowth [18] – two further processes controlled by cytoskeleton filaments. As a transmembrane protein KIAA0319 might have an involvement in linking the cellular cytoskeleton to the extracellular matrix. Further work should therefore expand our studies on surfaces coated with specific extracellular matrix proteins. Due to its involvement in the regulation of cilia and cell forces, we speculate that the *KIAA0319* gene might play an important role during neurodevelopment. Such processes are being increasingly associated with neurodevelopmental disorders including schizophrenia, depression, bipolar disorder [41] and autism [42].

## Materials and Methods

### Cell culture

hTERT-RPE1 cells were generated by transfection with pGRN145, which expresses hTERT under the control of the MPSV promoter, and were kindly supplied by Dr. Andrea Bodnar, Geron Inc. Cell lines were cultured in complete media (DMEM F12 with 10% of fetal bovine serum and 1% Penicillin/Streptomycin), or in serum-free media (DMEM F12 with 1% Penicillin/Streptomycin) at 37 °C and 5% CO_2_.

### Plasmids

pSPgRNA was a gift from Charles Gersbach (Addgene plasmid #47108) [43]. pSPCas9n-2A-GFP (pSpCas9n(BB)-2A-GFP (PX461)) was a gift from Feng Zhang (Addgene plasmid #48140) [44]. KIAA0319-GFP was a gift from Antonio Velayos-Baeza [15].

### Cloning and transfection

KIAA0319 knockout cell lines were generated through a CRISPR-Cas9 double nicking strategy designed with the web-based tool developed by Hsu and collaborators (http://crispr.mit.edu) [45]. This strategy uses Cas9 nickase (Cas9n), a modified Cas9 in which one of the nuclease domains has been mutated, lowering the rate of off-target effects compared to Cas9 [44]. RPE1 cells were transfected with pSpCas9n(BB)-2A-GFP (PX461) and paired gRNAs, using Lipofectamine3000 (ThermoFisher). gRNAs were generated by cloning annealed oligonucleotides containing the protospacer sequence into the chimeric gRNA sequence in pSPgRNA linearised with BbsI, downstream of a U6 promoter (Table S1). Sequences targeted were AGCCACCCCACAGACTACCA and TAAATTCCATTCATAGTTGT on KIAA0319 exon 6. pSpCas9n(BB)-2A-GFP (PX461) contains a GFP expression cassette that acts as indicator of positive transfection. Twenty-four hours after transfection, 384 individual GFP positive cells (four 96 well plates) were isolated using Fluorescence Activated Cell Sorting (FACS) and plated onto 96 well plates coated with Poly-D-Lysine for clonal expansion.

### Screening

Fifty cells were successfully expanded for further analysis. PCR was performed in all clones using primers int6-7R and int5-6F, that amplify a 1311 sequence DNA flanking the site targeted with the gRNAs (Table S1). Amplicons were digested with the restriction enzyme StyI. One of the used gRNAs targets this sequence, hence mutations caused by this gRNA are likely to eliminate this site. Amplicons from the 7 clones that showed loss of a StyI site upon digestion were cloned into Zero Blunt TOPO (ThermoFisher K280020) and sequenced using primers SP6 and T7. We identified one of these lines as a homozygous knockout as it contains two types of deletions causing frameshifts and premature stop codons.

### Immunofluorescence

Cells on the ERISM micro-cavity were fixed with 4% paraformaldehyde (PFA) in PBS at room temperature for 20 minutes. Immediately after fixation, cells were permeabilised with 0.1%Triton X-100 for 3 minutes and blocked for 30 minutes with 1% BSA in PBS. Cells were then stained for vinculin using anti-vinculin antibody (Merck Millipore, cat. no. 90227, 1:250 in BSA solution, 1 hour at room temperature) and for actin using TRITC-conjugated phalloidin (MerckMillipore, cat. no. 90228, 1:500 in BSA solution, 1 hour at room temperature). The nuclei of the cells were stained with DAPI (MerckMillipore, 1:1,000 in BSA), at room temperature for 3 minutes.

RPE1 cells for cilia analysis were cultured on uncoated coverslips for 48 hours with serum-free media, fixed with 4% PFA for 10 minutes, permeabilised with 0.1%Triton X-100, blocked with 10% goat serum in PBS, and stained with the ciliary marker ARL13B Antibody Rabbit polyclonal (17711-1-AP Proteintech) and anti-gamma-tubulin (Abcam 11316). Under serum starvation, cells stay in G_0_ and form cilia. We measured the length of the cilia manually using ImageJ. To ensure that cilia were positioned flat against the surface of the cell, only cilia that were completely in focus were considered.

### Gene expression quantification

qRT-PCRs were performed using Luna OneStep reagent (NEB) on biological triplicates. KIAA0319 expression was assessed with primers ex11F and ex12R (Table S1). Analysis was performed by the ΔΔCt method using Beta-actin as endogenous control. Results were normalised against expression in WT cells. Error bars are calculated using the standard deviation of the triplicates (2^ΔΔCt-s.d^ − 2^ΔΔCt+-s.d^).

### Western blot

Protein lysates were obtained from all cell lines using RIPA buffer and separated in a NuPAGE Bis-Tris 4-12% gradient gel (ThermoFisher). Proteins were transferred to a nitrocellulose membrane, blocked in WesternBreeze blocker (ThermoFisher) and incubated with primary antibodies anti-GFP (Chromotek #029762) and anti-beta actin (Sigma). Secondary antibodies were donkey anti-rat and anti-mouse HRP conjugated. Membranes were developed using SuperSignal WestFemto substrate (ThermoFisher).

### Scratch assay

The scratch assay is a simple way to measure cell migration in vitro and consists on creating a “scratch” on a confluent layer of cells and quantifying the movement of the cells over time to close this gap [46]. Since this test is performed in serum free culture conditions, which prevent the cells from dividing, it only takes into account cell movement and not proliferation. Wild type and Ex6KO cell lines were plated on a 6-well plastic plate (Nunclon Delta Surface,ThermoFisher Scientific). When confluent, the layer of cells was scratched with a pipette tip creating a straight gap. Cells were then washed with PBS to remove media and floating cells, and serum free media was added. We took images covering the whole gap at the time of the scratch (time 0) and after 24 hours. We measured the width of the scratch using TScratch [47], and calculated the mean width for each cell line after 24 hours.

### ERISM measurements

ERISM substrates were fabricated as described previously (Kronenberg, 2017) and four silicon chambers (surface area: 0.75 x 0.75 cm^2^; Ibidi) were applied. RPE1 cells were seeded on the ERISM substrate at 1,000 cells per well and kept at 37 °C, 5% CO_2_ culture conditions in DMEM-12 supplemented with 10% FBS and 1% Penicillin/Streptomycin. WT and Ex6KO cells as well as WT, WT_K-GFP, Ex6KO and Ex6KO_K-GFP cells were investigated in different wells on the same ERISM chip. Prior to ERISM measurements, cells were cultured for 24 h to allow adhesion to complete. ERISM force measurements were performed and converted into displacement maps as described before [21]. To investigate forces during cell migration, ERISM maps were recorded continuously for 17 h at intervals of 5 minutes, recording from seven different positions within each of the respective wells with a x20 objective. To analyse the force exertion patterns, ERISM measurements were performed at higher frame rate (every 5 s or 2 min) and magnification (x40 objective). To generate the Fourier-filtered ERISM maps, a FFT bandpass filter was applied to the raw displacement maps using the ImageJ software. For cell force analysis, the volume by which migrating cells indent into the ERISM substrate was calculated using ImageJ. All pixels in the ERISM displacement maps with indentation of less than 20 nm were set to NaN’s (not a number) and the “indented volume” under each individual cell was calculated as the product of area and mean indentation. Only cells that moved freely for >4 h (i.e. that were not in physical contact with other cells) were included in the analysis.

The “indentation force” of a single podosome protrusion was calculated by converting spatial Fourier-filtered ERISM displacement maps with a cut-off frequency of 0.6 µm^-1^ into stress maps using FEM as described in [21]. Podosome protrusions were identified in stress maps as isolated, localised indentation surrounded by a ring of pulling. Indentation force was calculated as the product of indentation area and mean applied stress at a threshold of 4 Pa. Only structures that colocalise with actin-dots in the respective immunostaining image were analysed.

To calculate the “contraction force” of single focal adhesions, the twist in spatial Fourier-filtered ERISM displacement maps with a cut-off frequency of 0.6 µm^-1^ were analysed and converted into the corresponding horizontally exerted contractile forces as described in [21]. In short, twisting results from the torque applied by focal adhesions when transmitting contractile cell forces to the ERISM substrate. The twisting response of ERISM substrates was calibrated by applying horizontal forces using AFM. The amount of twisting was found to be directly proportional to the applied force (6.6 nm of twist per 1 nN of applied force; *R*^2^ > 0.99; *n* = 5 force measurements). Only twists in ERISM displacement maps that form around vinculin-rich areas in the respective immunostaining image were analysed.

The speed of the cells on the ERISM sensor was measured using the plugin Manual Tracking on ImageJ[48]. The “straightness” of cell migration was calculated as the ratio of the effective displacement of the cell relative to the position at the start of the measurement and the track length. Cell areas were measured from single phase contrast images by drawing the outline of the cells in ImageJ..

### Generation of cell lines expressing KIAA0319-GFP

RPE1 wild type and Ex6KO were transfected with linearised KIAA0319-GFP plasmid using Lipofectamine 3000 according to the manufacturer’s specifications. KIAA0319-GFP contains a neomycin resistance cassette that was used to select cells that had undergone stable transfection, integrating the construct in their genome. Stably transfected cells were selected with G418 (Roche) at a concentration of 400 µg ml^-1^ for 2 weeks. Cells tend to lose the expression of the transgene with time [50], and after a few passages of this cell line, GFP expression was detected in only a small percentage of the cells. To enrich cells expressing the construct, we selected GFP positive cells via FACS. After FACS selection, cells were kept in culture for 24 hours to allow them to recover, and then plated onto the ERISM microcavity for measurement.

## Author contributions

RD and NMK conducted the experiments and analyzed the data with support from AM and PL. ACR assisted with the generation of the cell lines. RD and NMK wrote the manuscript with input from all authors. MCG and SP co-supervised the work.We thank Dr. Samantha Pitt and Dr. Swati Arya for their suggestions to improve the manuscript.

## Data and materials availability

All data needed to evaluate the conclusions in the paper are present in the paper and/or the Supplementary Materials. Additional data related to this paper are available via https://doi.org/….

## References

1. Peterson RL, Pennington BF. Developmental dyslexia. Lancet. 2012;379: 1997–2007. doi:10.1016/S0140-6736(12)60198-6

2. Paracchini S, Diaz R, Stein J. Advances in Dyslexia Genetics—New Insights Into the Role of Brain Asymmetries. Advances in Genetics. Elsevier Ltd; 2016. doi:10.1016/bs.adgen.2016.08.003

3. Paracchini S, Scerri T, Monaco AP. The genetic lexicon of dyslexia. Annu Rev Genomics Hum Genet. 2007;8: 57–79. doi:10.1146/annurev.genom.8.080706.092312

4. Galaburda AM, Sherman GF, Rosen GD, Aboitz F, Geschwind N. Developmental Dyslexia: Four consecutive patients with cortical anomaly. Ann Neurol. 1985;18: 222–223. doi:10.1002/ana.410180210

5. Guidi LG, Velayos-Baeza A, Martinez-Garay I, Monaco AP, Paracchini S, Bishop DVM, et al. The Neuronal Migration Hypothesis of Dyslexia: A Critical Evaluation Thirty Years On. Eur J Neurosci. 2018; 3212–3233. doi:10.1111/ejn.14149

6. Rosen GD, Bai J, Wang Y, Fiondella CG, Threlkeld SW, Loturco JJ, et al. Disruption of neuronal migration by RNAi of Dyx1c1 results in neocortical and hippocampal malformations. Cereb Cortex. 2007;17: 2562–2572. doi:10.1093/cercor/bhl162

7. Galaburda AM. The Role of Rodent Models in Dyslexia Research: Understanding the Brain, Sex Differences, Lateralization, and Behavior. In: Lachmann T, Weis T, editors. Reading and Dyslexia: From Basic Functions to Higher Order Cognition. Cham: Springer International Publishing; 2018. pp. 77–96. doi:10.1007/978-3-319-90805-2_5

8. Centanni TM. Neural and Genetic Mechanisms of Dyslexia. In: Argyropoulos GPD, editor. Translational Neuroscience of Speech and Language Disorders. Cham: Springer International Publishing; 2020. pp. 47–68. doi:10.1007/978-3-030-35687-3_4

9. Geremek M, Ziȩtkiewicz E, Bruinenberg M, Franke L, Pogorzelski A, Wijmenga C, et al. Ciliary genes are down-regulated in bronchial tissue of primary ciliary dyskinesia patients. PLoS One. 2014;9. doi:10.1371/journal.pone.0088216

10. Hoh R a, Stowe TR, Turk E, Stearns T. Transcriptional program of ciliated epithelial cells reveals new cilium and centrosome components and links to human disease. PLoS One. 2012;7: e52166. doi:10.1371/journal.pone.0052166

11. Ivliev AE, ’t Hoen P a C, WMC van Roon-Mom, Peters DJM, Sergeeva MG. Exploring the transcriptome of ciliated cells using in silico dissection of human tissues. PLoS One. 2012;7:e35618. doi:10.1371/journal.pone.0035618

12. Brandler WM, Paracchini S. The genetic relationship between handedness and neurodevelopmental disorders. Trends Mol Med. 2013; 1–8. doi:10.1016/j.molmed.2013.10.008

13. Pruski M, Lang B. Primary Cilia–An Underexplored Topic in Major Mental Illness. Front Psychiatry. 2019;10. doi:10.3389/FPSYT.2019.00104

14. Velayos-Baeza A, Toma C, da Roza S, Paracchini S, Monaco AP. Alternative splicing in the dyslexia-associated gene KIAA0319. Mamm Genome. 2007;18: 627–34. doi:10.1007/s00335-007-9051-3

15. Velayos-Baeza A, Toma C, Paracchini S, Monaco AP. The dyslexia-associated gene KIAA0319 encodes highly N- and O-glycosylated plasma membrane and secreted isoforms. Hum Mol Genet. 2008;17: 859–71. doi:10.1093/hmg/ddm358

16. Ibraghimov-Beskrovnaya O, Bukanov NO, Donohue LC, Dackowski WR, Klinger KW, Landes GM. Strong homophilic interactions of the Ig-like domains of polycystin-1, the protein product of an autosomal dominant polycystic kidney disease gene, PKD1. Hum Mol Genet. 2000;9: 1641–9. Available: http://www.ncbi.nlm.nih.gov/pubmed/10861291

17. Velayos-Baeza A, Levecque C, Kobayashi K, Holloway ZG, Monaco AP. The dyslexia-associated KIAA0319 protein undergoes proteolytic processing with {gamma}-secretase-independent intramembrane cleavage. J Biol Chem. 2010;285: 40148–62. doi:10.1074/jbc.M110.145961

18. Franquinho F, Nogueira-Rodrigues J, Duarte JM, Esteves SS, Carter-Su C, Monaco AP, et al. The Dyslexia-susceptibility Protein KIAA0319 Inhibits Axon Growth Through Smad2 Signaling. Cereb Cortex. 2017;27: 1732–1747. doi:10.1093/cercor/bhx023

19. Gostic M, Martinelli A, Tucker C, Yang Z, Ewart FGJ, Dholakia K, et al. The dyslexia susceptibility KIAA0319 gene shows a specific expression pattern during zebrafish development supporting a role beyond neuronal migration. J Comp Neurol. 2019; 1–10. doi:10.1002/cne.24696

20. Wu G De, Li ZH, Li X, Zheng T, Zhang DK. microRNA-592 blockade inhibits oxidative stress injury in Alzheimer’s disease astrocytes via the KIAA0319-mediated Keap1/Nrf2/ARE signaling pathway. Exp Neurol. 2020;324: 113128. doi:10.1016/j.expneurol.2019.113128

21. Kronenberg NM, Liehm P, Steude A, Knipper JA, Borger JG, Scarcelli G, et al. Long-term imaging of cellular forces with high precision by elastic resonator interference stress microscopy. Nat Cell Biol. 2017;19: 864–872. doi:10.1038/ncb3561

22. Liehm P, Kronenberg NM, Gather MC. Analysis of the Precision, Robustness, and Speed of Elastic Resonator Interference Stress Microscopy. Biophys J. 2018; 2180–2193. doi:10.1016/j.bpj.2018.03.034

23. Labernadie A, Bouissou A, Delobelle P, Balor S, Voituriez R, Proag A, et al. Protrusion force microscopy reveals oscillatory force generation and mechanosensing activity of human macrophage podosomes. Nat Commun. 2014;5: 5343. doi:10.1038/ncomms6343

24. Baker KE, Parker R. Nonsense-mediated mRNA decay: terminating erroneous gene expression. Curr Opin Cell Biol. 2004;16: 293–299. doi:10.1016/J.CEB.2004.03.003

25. Legant WR, Choi CK, Miller JS, Shao L, Gao L, Betzig E, et al. Multidimensional traction force microscopy reveals out-of-plane rotational moments about focal adhesions. Proc Natl Acad Sci. 2013;110: 1–6. doi:10.1073/pnas.1207997110

26. Linder S, Wiesner C. Feel the force: Podosomes in mechanosensing. Exp Cell Res. 2016;343: 67–72. doi:10.1016/J.YEXCR.2015.11.026

27. Spinardi L, Rietdorf J, Nitsch L, Bono M, Tacchetti C, Way M, et al. A dynamic podosome-like structure of epithelial cells. Exp Cell Res. 2004;295: 360–374. doi:10.1016/j.yexcr.2004.01.007

28. Kim S, Zaghloul NA, Bubenshchikova E, Oh EC, Rankin S, Katsanis N, et al. Nde1-mediated inhibition of ciliogenesis affects cell cycle re-entry. Nat Cell Biol. 2011;13: 351–362. doi:10.1038/ncb2183

29. Chandrasekar G, Vesterlund L, Hultenby K, Tapia-Páez I, Kere J. The zebrafish orthologue of the dyslexia candidate gene DYX1C1 is essential for cilia growth and function. PLoS One. 2013;8: e63123. doi:10.1371/journal.pone.0063123

30. Massinen S, Hokkanen M-E, Matsson H, Tammimies K, Tapia-Páez I, Dahlström-Heuser V, et al. Increased expression of the dyslexia candidate gene DCDC2 affects length and signaling of primary cilia in neurons. PLoS One. 2011;6: e20580. doi:10.1371/journal.pone.0020580

31. Skardal A, Mack D, Atala A, Soker S. Substrate elasticity controls cell proliferation, surface marker expression and motile phenotype in amniotic fluid-derived stem cells. J Mech Behav Biomed Mater. 2013;17: 307–316. doi:10.1016/j.jmbbm.2012.10.001

32. Bangasser BL, Shamsan GA, Chan CE, Opoku KN, Tüzel E, Schlichtmann BW, et al. Shifting the optimal stiffness for cell migration. Nat Commun. 2017;8: 15313. doi:10.1038/ncomms15313

33. Dalagiorgou G, Basdra EK, Papavassiliou AG. Polycystin-1: Function as a mechanosensor. Int J Biochem Cell Biol. 2010;42: 1610–1613. doi:10.1016/J.BIOCEL.2010.06.017

34. Qian F, Wei W, Germino G, Oberhauser A. The nanomechanics of polycystin-1 extracellular region. J Biol Chem. 2005;280: 40723–30. doi:10.1074/jbc.M509650200

35. Boca M, D’amato L, Distefano G, Polishchuk RS, Germino GG, Boletta A. Polycystin-1 Induces Cell Migration by Regulating Phosphatidylinositol 3-kinase-dependent Cytoskeletal Rearrangements and GSK3-dependent Cell-Cell Mechanical Adhesion. Mol Biol Cell. 2007;18: 4050–4061. doi:10.1091/mbc.E07-02-0142

36. Besschetnova TY, Kolpakova-Hart E, Guan Y, Zhou J, Olsen BR, Shah J V. Identification of Signaling Pathways Regulating Primary Cilium Length and Flow-Mediated Adaptation. Curr Biol. 2010;20: 182–187. doi:10.1016/J.CUB.2009.11.072

37. Hiroyasu S, Colburn ZT, Jones JCR. A hemidesmosomal protein regulates actin dynamics and traction forces in motile keratinocytes. FASEB J. 2016;30: 2298–2310. doi:10.1096/fj.201500160R

38. Zhang H, Landmann F, Zahreddine H, Rodriguez D, Koch M, Labouesse M. A tension-induced mechanotransduction pathway promotes epithelial morphogenesis. Nature. 2011;471: 99–103. doi:10.1038/nature09765

39. Grashoff C, Hoffman BD, Brenner MD, Zhou R, Parsons M, Yang MT, et al. Measuring mechanical tension across vinculin reveals regulation of focal adhesion dynamics. Nature. 2010;466: 263–266. doi:10.1038/nature09198

40. Walko G, Castañón MJ, Wiche G. Molecular architecture and function of the hemidesmosome. Cell Tissue Res. 2015;360: 363–378. doi:10.1007/s00441-014-2061-z

41. Marchisella F, Coffey ET, Hollos P. Microtubule and microtubule associated protein anomalies in psychiatric disease. Cytoskeleton. 2016;73: 596–611. doi:10.1002/cm.21300

42. Lin Y-C, Frei JA, Kilander MBC, Shen W, Blatt GJ. A Subset of Autism-Associated Genes Regulate the Structural Stability of Neurons. Front Cell Neurosci. 2016;10: 263. doi:10.3389/fncel.2016.00263

43. Perez-Pinera P, Kocak DD, Vockley CM, Adler AF, Kabadi AM, Polstein LR, et al. RNA-guided gene activation by CRISPR-Cas9 – based transcription factors. Nat Methods. 2013;10: 973–976. doi:10.1038/NMETH.2600

44. Ran FA, Hsu PD, Wright J, Agarwala V, Scott D a, Zhang F. Genome engineering using the CRISPR-Cas9 system. Nat Protoc. 2013;8: 2281–308. doi:10.1038/nprot.2013.143

45. Hsu PD, Scott DA, Weinstein JA, Ran FA, Konermann S, Agarwala V, et al. DNA targeting specificity of RNA-guided Cas9 nucleases. Nat Biotechnol. 2013;31: 827–832. doi:10.1038/nbt.2647

46. Liang CC, Park AY, Guan JL. In vitro scratch assay: a convenient and inexpensive method for analysis of cell migration in vitro. Nat Protoc. 2007;2: 329–333. doi:nprot.2007.30 [pii]\r10.1038/nprot.2007.30

47. Geback T, Schulz MMP, Koumoutsakos P, Detmar M. TScratch: A novel and simple software tool for automated analysis of monolayer wound healing assays. Biotechniques. 2009;46: 265– 274. doi:10.2144/000113083

48. Schneider CA, Rasband WS, Eliceiri KW. NIH Image to ImageJ: 25 years of image analysis. Nat Methods. 2012;9: 671–675.

49. Schindelin J, Arganda-Carreras I, Frise E, Kaynig V, Longair M, Pietzsch T, et al. Fiji: an open-source platform for biological-image analysis. Nat Methods. 2012;9: 676–682. doi:10.1038/nmeth.2019

50. Mutskov V, Felsenfeld G. Silencing of transgene transcription precedes methylation of promoter DNA and histone H3 lysine 9. EMBO J. 2004;23: 138–49. doi:10.1038/sj.emboj.7600013

51. Zerbino DR, Achuthan P, Akanni W, Ridwan Amode M, Barrell D, Bhai J, et al. Ensembl 2018. Nucleic Acids Res. 2018;46. doi:10.1093/nar/gkx1098

