## Supplementary Information for "KIAA0319 influences cilia length, cell migration and mechanical cell-substrate interaction"

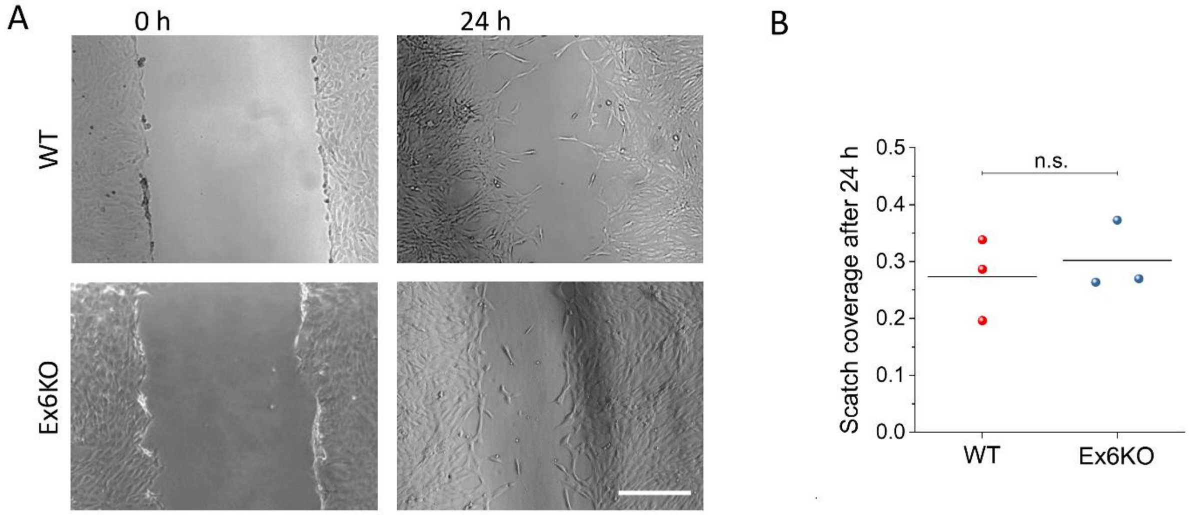

**Figure S1. Scratch assay**
**(A)** Representative phase contrast microscopy images of scratch assay at time 0 and after 24 h, for RPE1 wild type and Ex6KO. **(B)** Ratio of the scratch covered for each cell line after 24 h. Each dot represents an independent experiment ($n$ = 3). There is no significant difference between the cell lines ($t$-test, $p$ = 0.63). Scale bar, 500 µm.

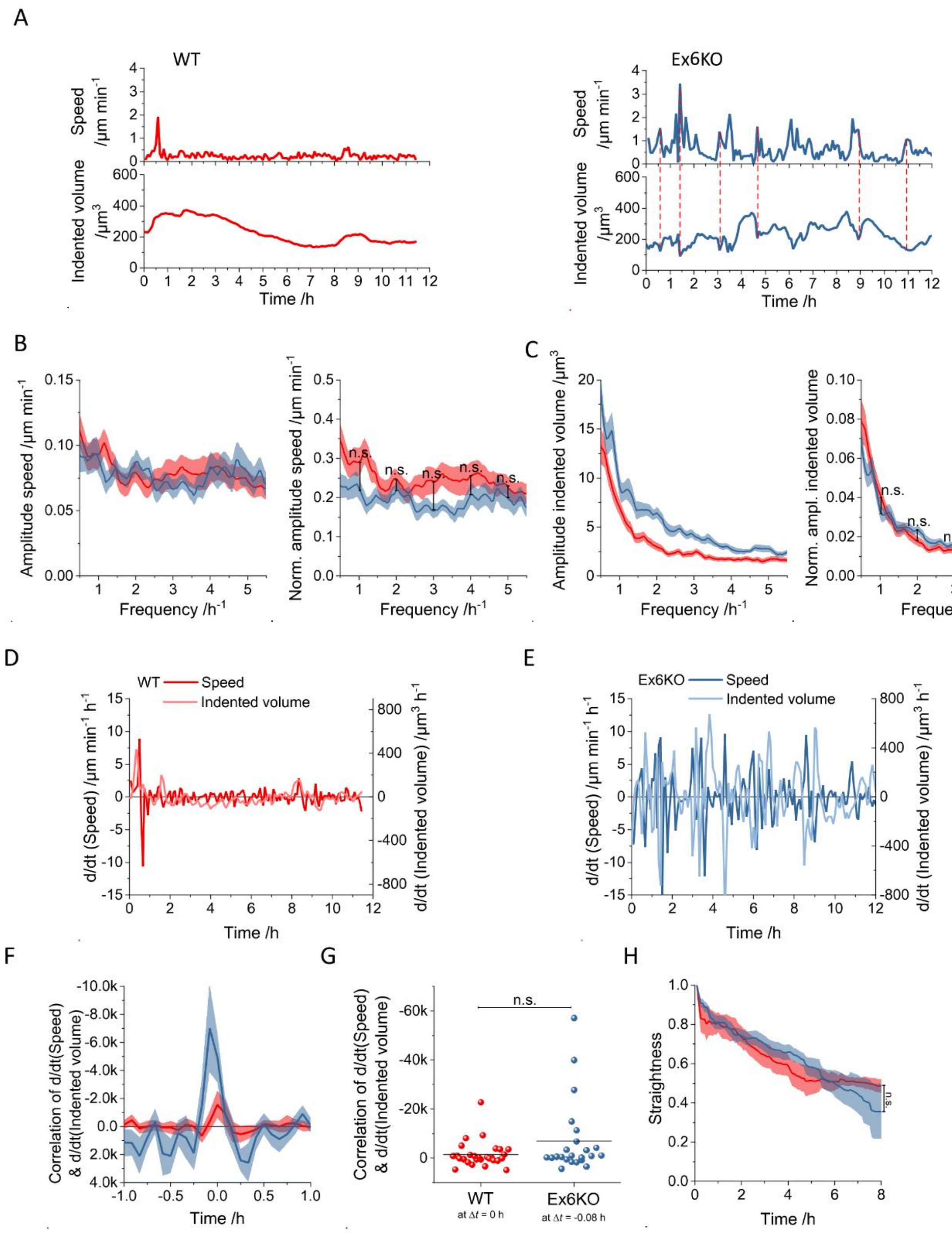

**Figure S2. Analysis of mechanical activity of RPE1 WT and Ex6KO cells during migration.** **(A)** Exemplary temporal evolution of speed and mechanical activity (using the total volume by which each cell indents into the ERISM substrate as a proxy for the applied force) of representative RPE1 WT (left panel) and Ex6KO (right panel) cells, following the movement of two individual cells on an ERISM micro-cavity for >11 h. Red, vertical lines indicate time points when high migration speed of Ex6KO cells correlate with a drop in exerted force. FFT analysis of amplitude of the oscillation of migration speed **(B)** and mechanical activity **(C)** of single RPE1 WT (red) and Ex6KO (blue) cells during migration on ERISM micro-cavity. For significance testing via *t*-test, the mean amplitudes of the individual cells were calculated separately for each 1 $h^{-1}$ frequency range. First time derivatives of speed and mechanical activity of **(D)** the WT cell and **(E)** the Ex6KO cell shown in (A). **(F)** Temporal correlation between the first time derivatives of speed and mechanical activity (see D & E) of single RPE1 WT (red) and Ex6KO (blue). **(G)** Comparison of the temporal correlation between the first time derivatives of speed and mechanical activity for WT ($n$ = 29) and Ex6KO ($n$ = 24) cells for $\Delta t$ = 0. Each data point represents the measurement for one cell. Only cells with free movement for >2 h were included in analysis. The lines depict the positions of the means. Groups were compared using the Student's $t$-test (*: $p \leq 0.05$). **(H)** Temporal evolution of straightness of RPE1 WT (red) and Ex6KO (blue) single cell movement on ERISM micro-cavity. $t$-test was performed on data distributions after 8 h. In the plots in (B), (C), (F) and (H), lines depict the means and the shaded areas the standard error of the mean. Only cells with free movement for ≥2 h were included in the analysis. WT: $n = 29$, Ex6KO: $n = 24$.

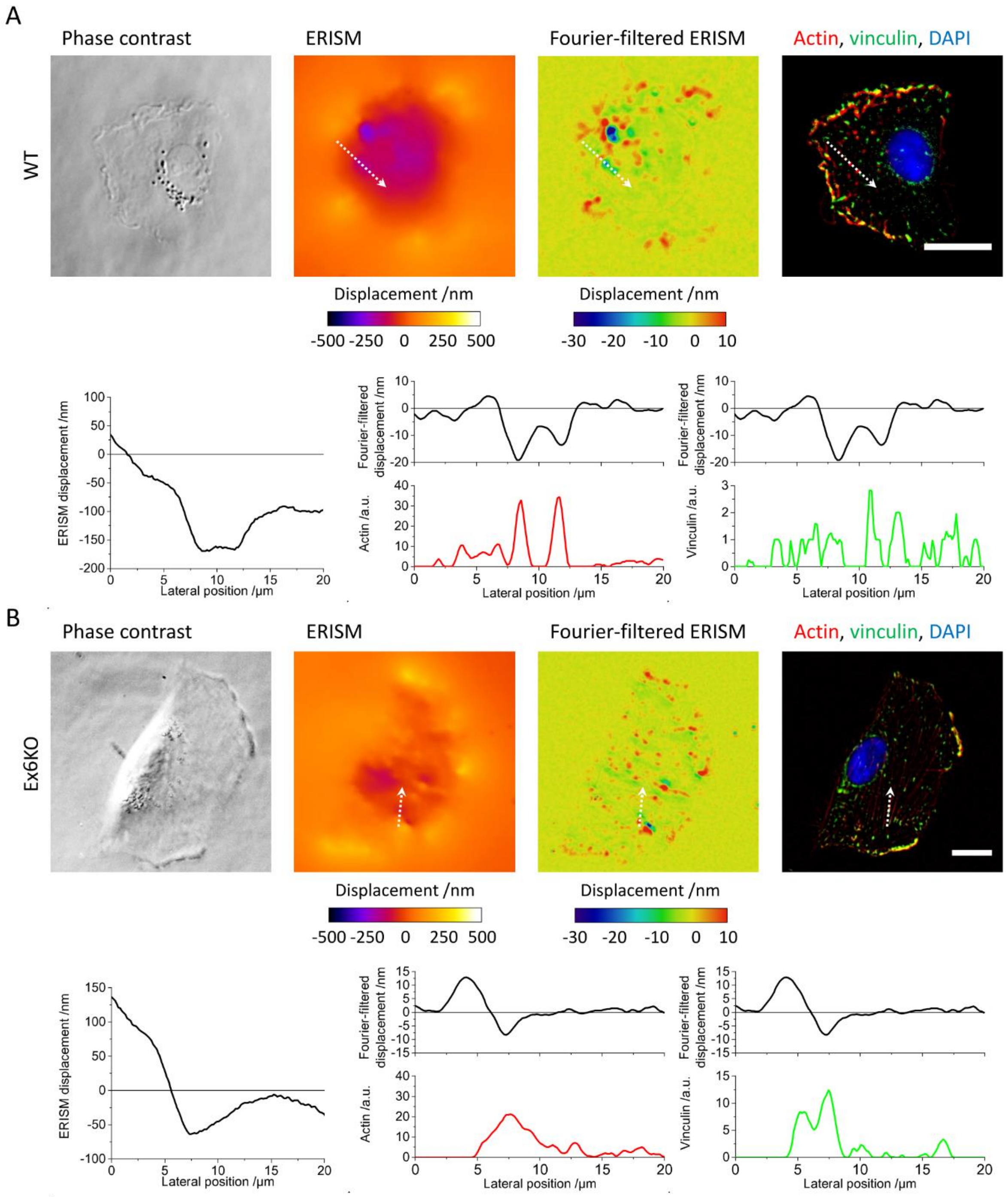

**Figure S3. Fourier-filtered ERISM displacement map reveals two different types of mechanical cell-substrate interaction in RPE1 cells**

Phase contrast images (upper row, left), ERISM displacement map (upper row, middle left), Fourier-filtered ERISM displacement maps (upper row, middle right) and epi-fluorescence images (upper row, right; red: actin, green: vinculin, blue: nuclear DNA) of WT **(A)** and Ex6KO **(B)** cell. The lower rows in A and B show the ERISM topography profile (left), the Fourier-filtered ERISM topography profile and the actin fluorescence intensity profile (middle), and the Fourier-filtered ERISM topography profile and the vinculin fluorescence intensity profile (right) measured along the line and direction indicated by dotted arrows in the respective images. All scale bars: 20 µm. Fourier-filtering of ERISM displacement maps reveals fine displacement features that are concealed in the unfiltered maps due to the overall cell contractility. The example of the WT cell in (A) shows tightly localised pushing sites in the Fourier-filtered ERISM map that colocalise with actin and are surrounded by rings of pulling sites that colocalise with vinculin, suggesting that the displacement is caused by podosome-associated actin and vinculin. The example of the Ex6KO cell in (B) shows a push-pull pattern in the Fourier-filtered ERISM displacement map that colocalises with vinculin expression and is aligned along the direction of an actin stress fibre, suggesting that this feature is related to the torque exerted by a focal adhesion transmitting actomyosin contraction.

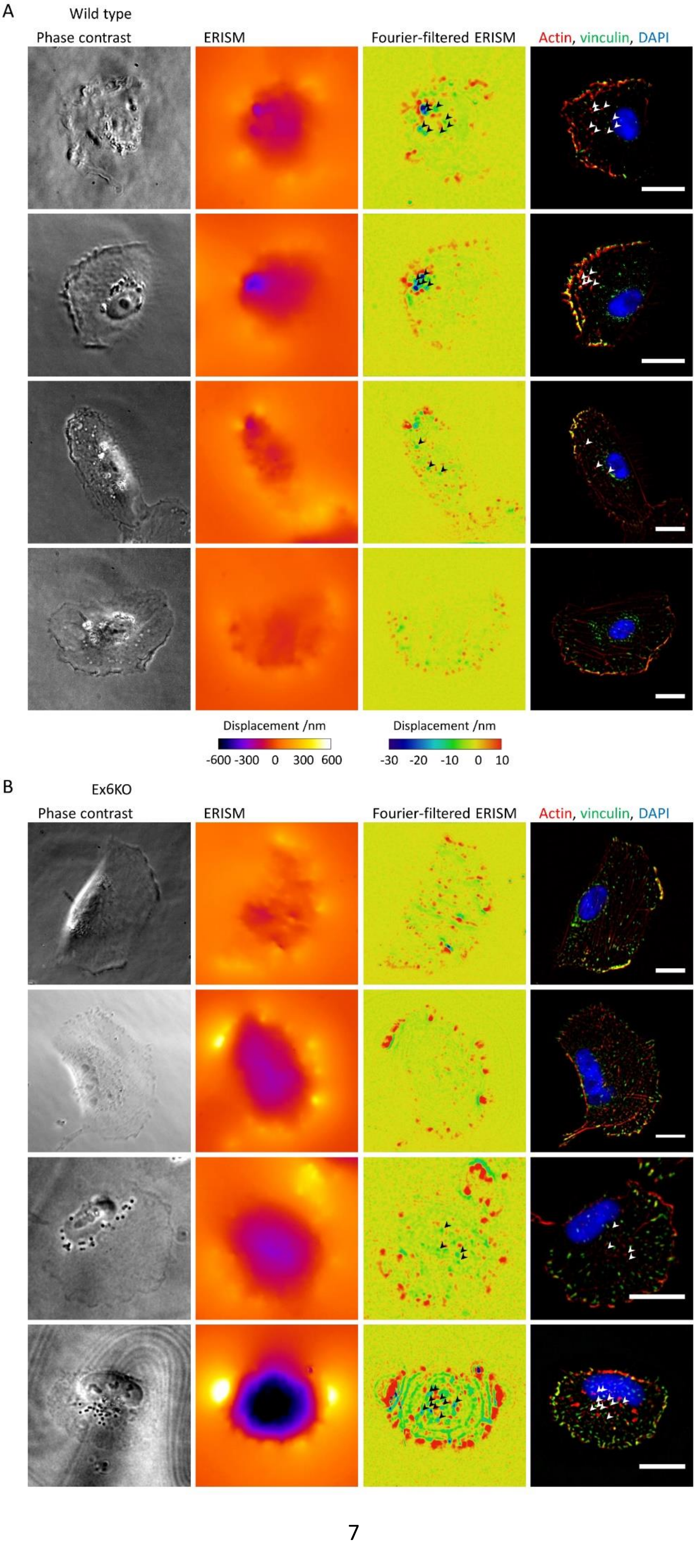
A
Wild type
Phase contrast
ERISM
Fourier-filtered ERISM
Actin, vinculin, DAPI
Displacement /nm
-600 -300 0 300 600
Displacement /nm
-30 -20 -10 0 10
B
Ex6KO
Phase contrast
ERISM
Fourier-filtered ERISM
Actin, vinculin, DAPI

66

**Figure S4. RPE1 KIAA0319 WT and Ex6KO cells use different modes of force exertion**

Phase contrast images (left columns), displacement maps (centre left columns), Fourier-filtered ERISM displacement maps (centre right columns) and epi-fluorescence images (right columns) of three fixed RPE1 WT cells **(A)** and three fixed Ex6KO cells **(B)** with staining for actin (red), vinculin (green) and nuclear DNA (blue). White circles in Fourier-filtered ERISM displacement maps and epi-fluorescence images indicate positions of vinculin-rich cell-substrate contacts (focal adhesions). White arrow heads in Fourier-filtered ERISM displacement maps and epi-fluorescence images indicate positions of actin-rich cell-substrate contacts (podosomes). All scale bars: 20 µm.

**Movie S1. ERISM time-lapse investigation of mechanical activity of RPE1 WT cells**
Time-lapse movie of phase contrast (left) and ERISM displacement (right) of RPE1 WT cells migrating on an ERISM substrate taken in intervals of five minutes over a time span of 17 hours.

**Movie S2. ERISM time-lapse investigation of mechanical activity of RPE1 Ex6KO cells**
Time-lapse movie of phase contrast (left) and ERISM displacement (right) of RPE1 Ex6KO cells migrating on an ERISM substrate taken in intervals of five minutes over a time span of 17 hours.

**Movie S3. ERISM time-lapse investigation of mechanical activity of an RPE1 WT cell**
Time-lapse movie of phase contrast (left) and Fourier-filtered ERISM displacement (right) of a RPE1 WT cell taken in intervals of five seconds over a time span of 12.5 minutes.

**Movie S4. ERISM time-lapse investigation of mechanical activity of an RPE1 Ex6KO cell**
Time-lapse movie of phase contrast (left) and Fourier-filtered ERISM displacement (right) of a RPE1 Ex6KO cell taken in intervals of five seconds over a time span of 12.5 minutes.

**Movie S5. ERISM time-lapse investigation of mechanical activity of RPE1 WT cells shown in Figure S4.**
Phase contrast image (left) and time-lapse movie of ERISM displacement (middle) and Fourier-filtered ERISM displacement (right) of three RPE1 WT cells taken in intervals of two minutes over a time span of 50 minutes. The displacement maps of the upper two cells show local, vertical force exertion by podosomes. The existence of podosomes in these two cells was later confirmed by immunostaining (see Figure S4).

**Movie S6. ERISM time-lapse investigation of mechanical activity of RPE1 Ex6KO cells shown in Figure S4.**
Phase contrast image (left) and time-lapse movie of ERISM displacement (middle) and Fourier-filtered ERISM displacement (right) of three RPE1 Ex6KO cells taken in intervals of two minutes over a time span of 42 minutes. The displacement maps of the lower two cells show local, vertical force exertion by podosomes. The existence of podosomes in these two cells was later confirmed by immunostaining (see Figure S4).

**Table S1. Primer sequences**

| | | use | Size (in bp) |
|---|---|---|---|
| int6-7R | ATCTAAGGTAATCTGCACTGGTGG | PCR | 1311 |
| int5-6F | AAATTAGCCGGGTGTGGTGAC | | |
| ex11F | TCTTCAAGGCAACAGTCTACTG | qRT-PCR | 128 |
| ex12R | CCATCCAGGGTAGCACTTTC | | |
| Ex6_R | AGAGTTTGCTTGTGTCCTTG | RT-PCR | 137 |
| Ex5_F | CCCGACAATGAAGTTGAACTG | | |
| ex9R | ACGGGAGAGTCAACTGAAGTC | RT-PCR | 360 |
| ex6delF | CAACTATGAATGGAATTTAATAAGCCACC | | |
| NHEJ gBR | AAACTGGTAGTCTGTGGGGTGGCTC | gRNA generation | |
| NHEJ gBF | CACCGAGCCACCCCACAGACTACCA | | |
| NHEJ gAR | AAACACAACTATGAATGGAATTTAC | | |
| NHEJ gAF | CACCGTAAATTCCATTCATAGTTGT | | |
